# Featherweight long read alignment using partitioned reference indexes

**DOI:** 10.1101/386847

**Authors:** Hasindu Gamaarachchi, Sri Parameswaran, Martin A. Smith

## Abstract

The advent of nanopore sequencing has realised portable genomic research and applications. However, state of the art long read aligners and large reference genomes are not compatible with most mobile computing devices due to their high memory requirements. We show how memory requirements can be reduced through parameter optimization and reference genome partitioning, but highlight the associated limitations and caveats of these approaches. We then demonstrate how these issues can be overcome through an appropriate merging technique. We extend the Minimap2 aligner and demonstrate that long read alignment to the human genome can be performed on a system with 2GB RAM with negligible impact on accuracy.

## Introduction

Long read sequencing has revolutionised genome research by facilitating the characterisation of large structural variations, repetitive regions, and de-novo assembly of whole genomes. Pacific Biosciences (PacBio) and Oxford Nanopore Technologies (ONT) are leading manufacturers that produce long read sequencers. In particular, ONT manufacture sequencers smaller than the size of a mobile phone that can nevertheless output more than 1TB of data in 48 hours. Such highly portable sequencers have realised the possibility of performing genome sequencing in the field. For instance, ONT’s MinION sequencer has been used for Ebola virus surveillance in New Guinea [1], mobile Zika virus surveillance in Brazil [2], and for experiments on the International space station [3].

The advent of highly portable DNA sequencers raises the need for local data processing on devices such as mobile phones, tablets and laptops. Facilitating genomic data analysis on mobile devices avoids the need for high speed internet connections and enables real-time genomic tests and experiments. For Nanopore sequencers, a pico-ampere ionic current signal is produced for each DNA read, which is subsequently converted to nucleotide bases via applied machine learning models. Until recently, a high performance workstation (Quad-core i7 or Xeon processor, 16GB RAM, 1TB SSD) was required for live base calling, the process of converting ionic signal to nucleotide sequences.

Most genomic analyses depend on base calling as an initial step, which can be efficiently performed through GPGPU software implementations on graphics cards or, quite conveniently, on dedicated portable hardware (ONT manufacture one such device, termed ‘MinIT’). Next, base called reads are typically aligned to a reference, in case of reference guided assembly, or aligned to themselves in case of de-novo assembly. Subsequent analyses (i.e. consensus sequence generation, variant calling, methylation detection, etc) should follow this alignment step. Therefore, an alignment tool that can run on portable devices such as mobile phones, tablets and laptops is the next step in realising the full portability of the whole Nanopore processing pipeline.

Minimap2 [4] is a general purpose mapper that is compatible with both DNA and RNA sequences. It can align both long reads and short reads, either to a reference or an assembly contig. It first employs hashing followed by chaining for coarse grain alignment. Then it performs an optional base level alignment using an optimised implementation of the Suzuki-Kasahara DP formulation [5]. Minimap2 stands out as the current aligner of choice for long reads, among other long read aligners such as BLASR [6], GraphMap [7], Kart [8], NGMLR [9] and LAMSA [10]; not only is it 30 times faster than existing long read aligners, but its accuracy is on par or superior to other algorithms [4]. Hash table based approach in Minimap2 has shown to be effective against long reads. In contrast, FM-index [11] based short read aligners such as BWA [12] and Bowtie [13] have shown to fail with ultra long reads (i.e. several hundred kilo bases or more) [14].

Most alignment tools build an index of the reference sequence (suffix tree, hash map, FM-index, etc) and store in volatile memory. While this is manageable for small genomes such as individual bacteria (~5Mbases), fungi (~50Mbases) or insects (~400Mbases), most vertebrate and some plant species require large amounts of memory (genomes in the 1-100 Gbase range). Here, we describe an approach for long read alignment to large reference genomes (or collections of genomes) with Minimap2 by splitting the index into smaller partitions. We expose caveats of this approach and describe how to circumvent them. We also compare the accuracy of the alignments from our partitioned index to a standard workflow using simulated long reads, Nanopore reference human genome sequencing data, and a 470kb long chromothriptic read from a human cancer cell line.

## Results

### Effect of parameters on the memory usage

With default options, Minimap2 requires more than 11GB of memory to align Nanopore reads to a human genome (See Supplementary Table S1). Considering the memory requirements of the operating system and other software, a computer with at least 16GB of memory is required to efficiently run Minimap2. Hence, running Minimap2 with default options on a typical laptop with 8GB memory or a typical mobile phone with 2GB of memory is not possible.

We therefore tested the effect of alignment parameters on peak memory usage in Minimap2 (see Materials and Methods), including the minimiser k-mer length (*k*: default 15 for Nanopore), minimiser window size (*w*: default 10), the number of threads (*t*: default 4) and the number of query bases loaded into memory at a time (*K*: default 500M). Parameters *k* and *w* considerably affect the peak memory usage for holding the index in memory (see Fig. 1a). For an index without homo-polymer compression, *k* = 15 consumed the least amount of memory out of the values tested. In fact, the default k-mer size for Oxford Nanopore in Minimap2 (set by *map-ont* pre-set) is 15.

**Figure 1:**
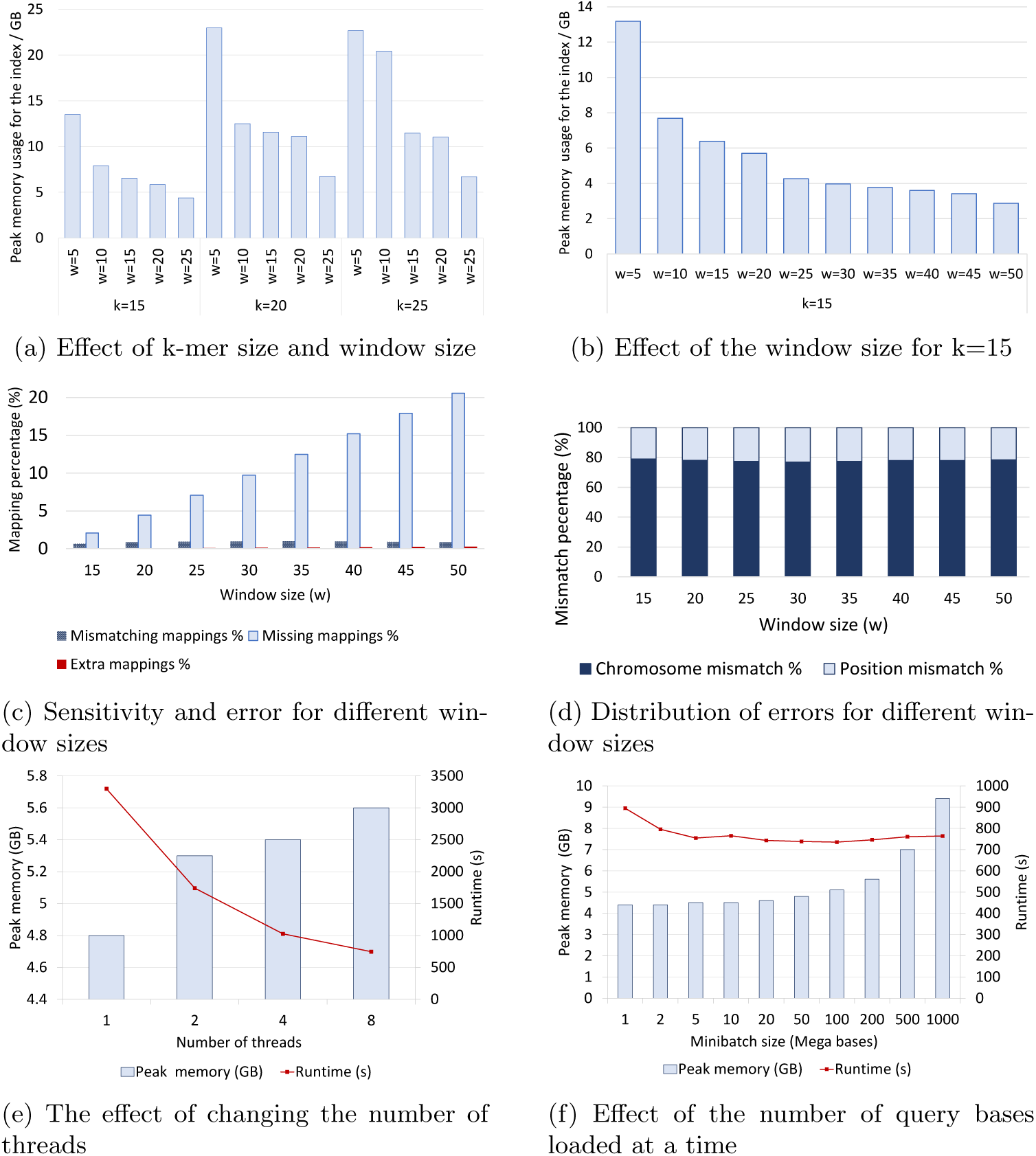
Effect of parameters on memory usage, performance and accuracy. (A) Peak memory usage of the index for different combinations of *k* and *w*. (**B**) Peak memory usage of the index for a large range of *w* with *k* =15. (**C**) The effect of *w* on sensitivity and error. The x-axis is the window size. The k-mer size is held constant at 15 for all values of *w*. The y-axis shows the number of missing mappings / mismatches or extra mapping as a percentage of the number of mappings for default *w* =10. (**D**) Error distribution for different window sizes. The mismatches in C are further classified. About 80% of the mismatches are chromosome mismatches and the remaining 20% are different loci in the same chromosome. (**E**) Effect of the number of threads on memory and performance. The parameters *k*, *w* and *K* were held constant at 15, 25 and 200M respectively while changing the number of threads. Both the peak memory usage and the runtime were measured on a PC with an Intel i7-6700 CPU and 16GB of RAM. (**F**) Effect of the number of query bases loaded at a time. The parameters *k*, *w* and *t* were held constant at 15, 25 and 8 respectively.

Memory usage decreases considerably when increasing *w* (see Fig. 1b), but this effect is mitigated with larger values of parameter *w*. At window size 50, memory usage is capped at 3GB. Unfortunately, increasing the window size heavily reduces the sensitivity (see Fig. 1c and materials and methods). A larger window size of 50 reduces sensitivity compared to the default window size by 20%, while a window size of 25 entails a reduction in sensitivity of about 7%. However, even a window size of 25 requires more than 4GB of memory. While sufficient for a computer with 8GB of RAM, this is still too high for smaller devices, such as mobile phones or microcomputer boards. Importantly, the error rate of mapped reads is not significantly affected by the window size parameter (see mismatches and extra bars in Fig. 1c). The number of threads marginally increases peak memory usage (see Fig. 1e). Only about 0.8GB of additional memory was consumed when moving from 1 to 8 threads, while producing a 6-fold gain in speed. Hence, reducing the number of threads for the sake of reduced memory usage is not an efficient solution.

Intuitively, the number of query bases loaded to the memory at once (also known as the mini-batch size) heavily impacts peak memory usage. This affects the size of the internal data structures used for mapping, but does not affect the index size. Hence, this parameter does not affect alignment accuracy or the sensitivity. A lower mini-batch size reduces the peak memory usage, at the cost of reduced multi-threading efficiency (see Fig. 1f). The runtime drops when changing *k* from 1M to 5M. However, the runtime is relatively stable from 5M onwards. It is important to note that the values in see Fig. 1f are only valid for 8 threads. A large number of threads would require a large mini-batch size for optimal performance.

Although a small window size value and mini-batch sizes from 5M-20M are suitable for systems with limited RAM (and 8 or lesser threads), tuning parameters alone cannot bring down the memory usage to a value lesser than 4GB. As a consequence, this inspired us to investigate the use and suitability of partitioned indexes.

### Caveats of naive partitioned indexes

Minimap2 allows the reference index to be split by a user specified number of bases through the option *I*, effectively dividing a reference into smaller indexes of comparable size. This facilitates parallel computation and, in theory, enables lower peak memory requirements. However, this feature is not ideal for mapping single reads to large references, mainly because global contiguous information about the reference is unavailable. As a result, several mapping artefacts can occur, as listed below and in Fig. 2 (N.B. these may not be as prominent when overlapping reads–the application for which index partitioning was originally developed).

**Figure 2:**
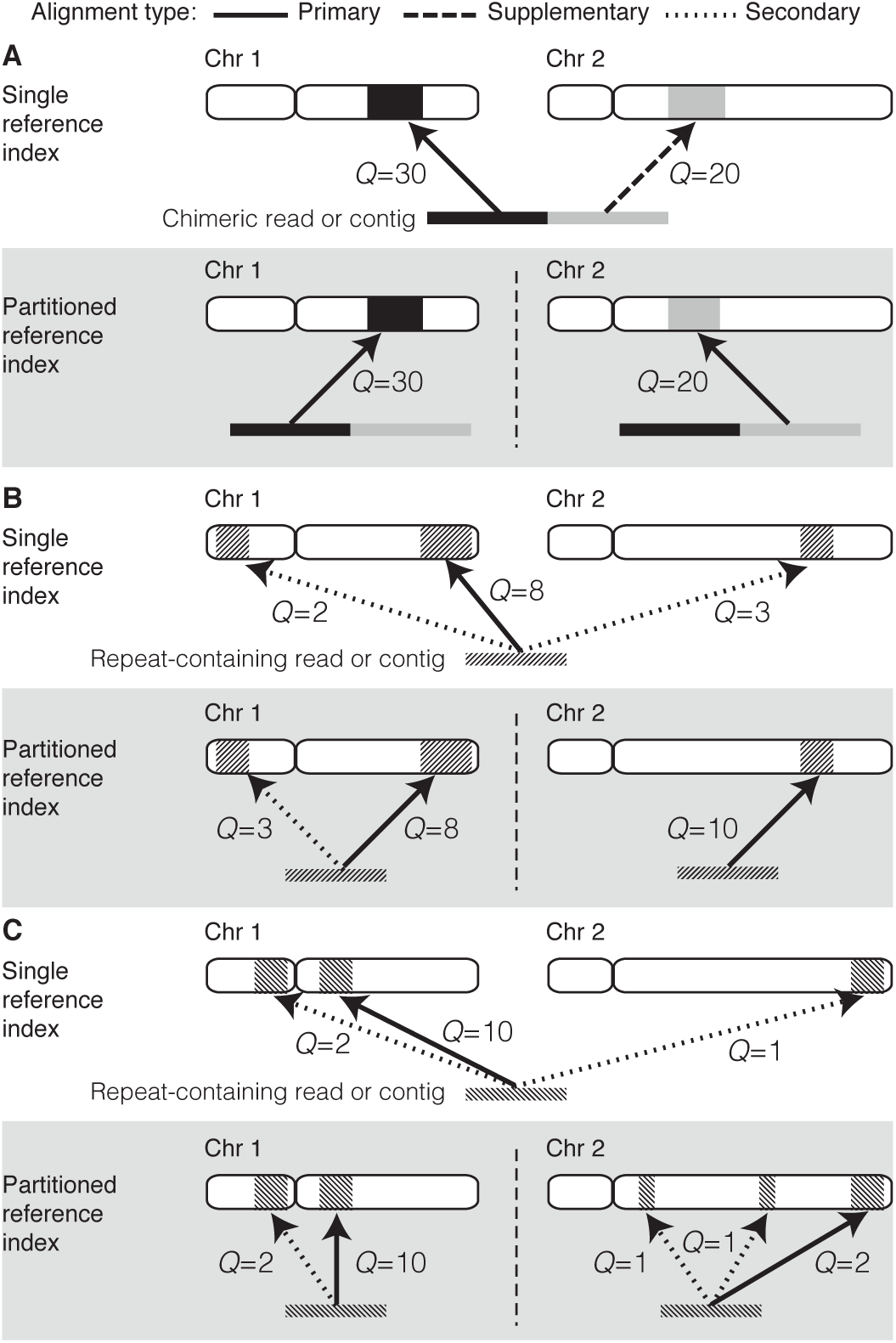
Effect of aligning sequences to single vs partitioned indexes. Uniquely mapping chimeric reads (**A**) can be reconstructed from a partitioned reference index with relative ease. However, sequences (or sub-sequences) that are difficult to map (i.e. low complexity regions, repetitive elements, etc) can cause artefacts when aligning to a partitioned reference index. (**B**) An example where one partition (chr2) contains less homologous sequences to the query sequence, producing the situation where the best alignment when using a single reference is not achieved. (**C**) An example where a partitioned reference introduces several additional low quality mappings that would be dismissed with a single reference index.

1. The mapping quality is incorrect. The mapping quality estimated in Minimap2 is accurate as it deliberately lowers the mapping quality for repetitive hits (see supplementary materials of [4]). However, this is not possible when only a fraction of a whole genome is present in the index. In a partitioned index, if the same repeat lies across different partitions, the mapping quality will be overestimated (fig2b.)
2. Incorrect alignment flags. For a chimeric read where different sub-sequences map to different chromosomes, the supplementary mappings would be marked as primary mappings (See fig2a). A repeat containing read that maps to multiple locations across different partitions will have multiple primary alignments instead of a single primary alignment (See fig2b and fig2c).
3. Large output files. For each partition of the index where a particular read does not map to, an spurious unmapped record will be printed. Furthermore, if a maximal amount of secondary alignments are specified, that number of secondary alignments would be output for each partition (See fig2c). Hence, the more partitions used, the larger the output files will be. Such large outputs not only waste disk space, but they are also time consuming to parse or sort.
4. Multiple hits of the same query may not be adjacent in the output [15] This causes difficulties to analyse or evaluate mapping results. For instance, the *Mapeval* utility in *Paftools* (a tool bundled with Minimap2 for evaluating alignment accuracy) is not compatible with such outputs. Sorting by the read identifier would fix the issue, but requires significant computations for large files.
5. Incomplete headers in the sequence alignment/map (SAM) output For a partitioned index, Minimap2 suppresses the reference sequence dictionary (SQ lines) in the SAM header. Users must manually add SQ lines to the header for compatibility with downstream analysis tools.

We resolved these issues by dumping the internal state of Minimap2 while mapping reads, then merging the output and processing the result *a posteriori* (see Materials and methods). The accuracy of this technique is discussed below.

### Effect of using a partitioned index on alignment accuracy

We compared the alignment accuracy between a single reference index and a partitioned index, with and without merging the output. The following acronyms will be used in the subsequent text (see Materials and methods for more details):

- *single-idx*: Aligning reads to a single reference index;
- *part-idx-no-merge*: Aligning reads to a partitioned index without merging the output;
- *part-idx-merged*: Aligning reads to a partitioned index while applying our merging technique.

## Synthetic long reads

Synthetic long reads were used as a ground truth for the evaluation of alignment accuracy (See Materials and Methods). The accuracy of *part-idx-merged* is similar to *single-idx*, despite employing significantly more partitions (fig3a and 3b) as exemplified by the overlap of their respective curves. In contrast, the results of *part-idx-no-merge* are considerably less accurate, in particular for larger quantities of index partitions. A lower error rate is observed for *part-idx-merged* when compared to *single-idx* for the lowest mapping quality values, but this effect is marginal and it is associated with low sensitivity.

**Figure 3:**
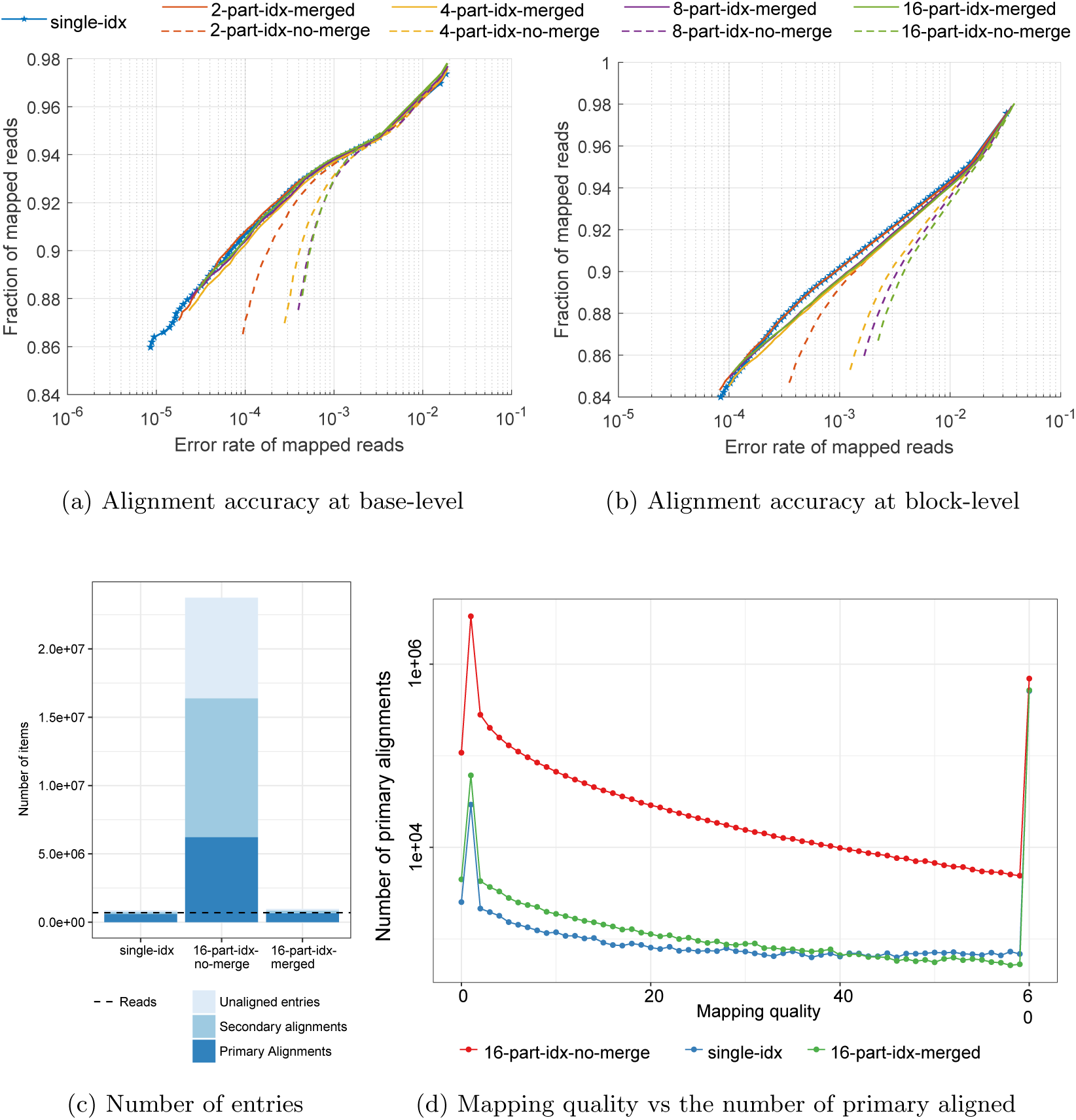
Effect of using partitioned indexes versus a single reference index on alignment quality Base-level. (**A**) and locus- or block-level (**B**) alignment accuracy from synthetic long reads. The x-axis shows the error rate alignments (see Materials and Methods). The y-axis shows the fraction of aligned reads out of all input reads. Each point in the plot corresponds to a mapping quality threshold that varies from 0 (top right) to 60 (bottom left). (**C**) and (**D**): Alignment statistics for real Nanopore whole genome sequencing data from [14] using a 16-part index. (**C**) The number of total entries (primary+secondary+unaligned) for *single-idx*, *16-part-idx-merged*, and *16-part-idx-no-merge*, in log scale. The dotted horizontal line represents the number of reads. (D) Number of primary mappings in function of Minimap2 mapping quality (log scale).

### For real Nanopore NA12878 reads

As no ground truth is available for real biological data, we evaluated the alignment accuracy by comparing a few alignment statistics such as the number of primary/secondary alignments and unmapped reads. When using *Single-idx*, Minimap2 ouputs 12.1GB of base-level alignment data, in SAM format. However, *part-idx-no-merge* generates much larger output (180GB), whereas *part-idx-merged* generates 12.4GB of data–almost identical to the output of *single-idx*. Hence, *part-idx-merged* reduces disk usage by about 14-fold compared to *part-idx-no-merge*. Peak disk usage is also minimised in *part-idx-merged* as only the alignments are dumped as temporary binary files. The resulting size of temporary files generated with *part-idx-merged* is 29.2GB, thus achieving maximal disk usage of 41.6GB (4 times less than *part-idx-no-merge*). The increased output in *part-idx-no-merge* is due to redundant unmapped entries and spurious mappings (fig3c and supplementary Table S2).

**Table S2:** Statistics for outputs for NA12878

|  | single-idx | 16-part-idx-no-merge | 16-part-idx-merged |
| --- | --- | --- | --- |
| **File size (SAM file)** | 12.1 GB | 180 GB | 12.4 GB |
| **No of reads** | 689 781 |  |  |
| **No of SAM entries** | 862 427 | 23 749 310 | 969 223 |
| **No of unaligned entries** | 127 177 | 7 365 979 | 120 775 |
| **No of aligned entries** | 735 250 | 16 383 331 | 848 448 |
| **No of primary alignments** | 592 748 | 6 228 567 | 654 302 |
| **No of secondary alignments** | 142 502 | 10 154 764 | 194 146 |

The number of total entries (primary+secondary+unaligned) for *single-idx* and *part-idx-merged* are comparable to the number of input reads (689,781), while *part-idx-no-merge* generates abundant (and presumably spurious) hits. Furthermore, the distribution of mapping qualities for primary alignments between *part-idx-merged* and *single-idx* are quite similar (fig3d). Interestingly, *part-idx-merged* produces slightly more primary alignments with lower mapping quality scores than *single-idx*, a likely consequence of sampling less repetitive regions in partitioned indexes. Both these strategies produce almost the same amount of mapping with quality = 60. In contrast, *part-idx-no-merge* has a very high number of spurious mappings for mapping qualities between 0 to 59.

### For an ultra-long chromothriptic read

To evaluate how chimeric reads will be affected by aligning them to multipart indexes, we tested this case on an ultra-long (473kb) chromothriptic nanopore read from a patient-derived liposarcoma cell line [16]. Chromothripsis is a genetic phenomenon often associated with cancer and congenital diseases. It is caused by several rounds of breakage-fusion-bridge, which produce complex and localised genomic rearrangements in a relatively short segment of DNA. The *single-idx* produced 41 (36 primary + 5 secondary mappings) mappings (fig4a). However, *part-idx-no-merge* (16 partitions) produced 688 (608 primary + 80 secondary) mappings (fig4b), while mapping with *part-idx-merged* resulted in 47 (42 primary + 5 secondary) mappings (fig4c).

In *single-idx* and *part-idx-merged*, 34 mappings were the same. Interestingly, there were 7 mappings unique to *single-idx* and 6 unique to *part-idx-merged* (See supplementary Fig. S1). All 7 alignments unique to *single-idx* map to the centromeric region of the chromosome 11 (4a), which is composed of large arrays of repetitive DNA (also known as satellite DNA). The alignments that are unique to *part-idx-merged* map to simple repeats (e.g. GAGAGAGA).

**Figure 4:**
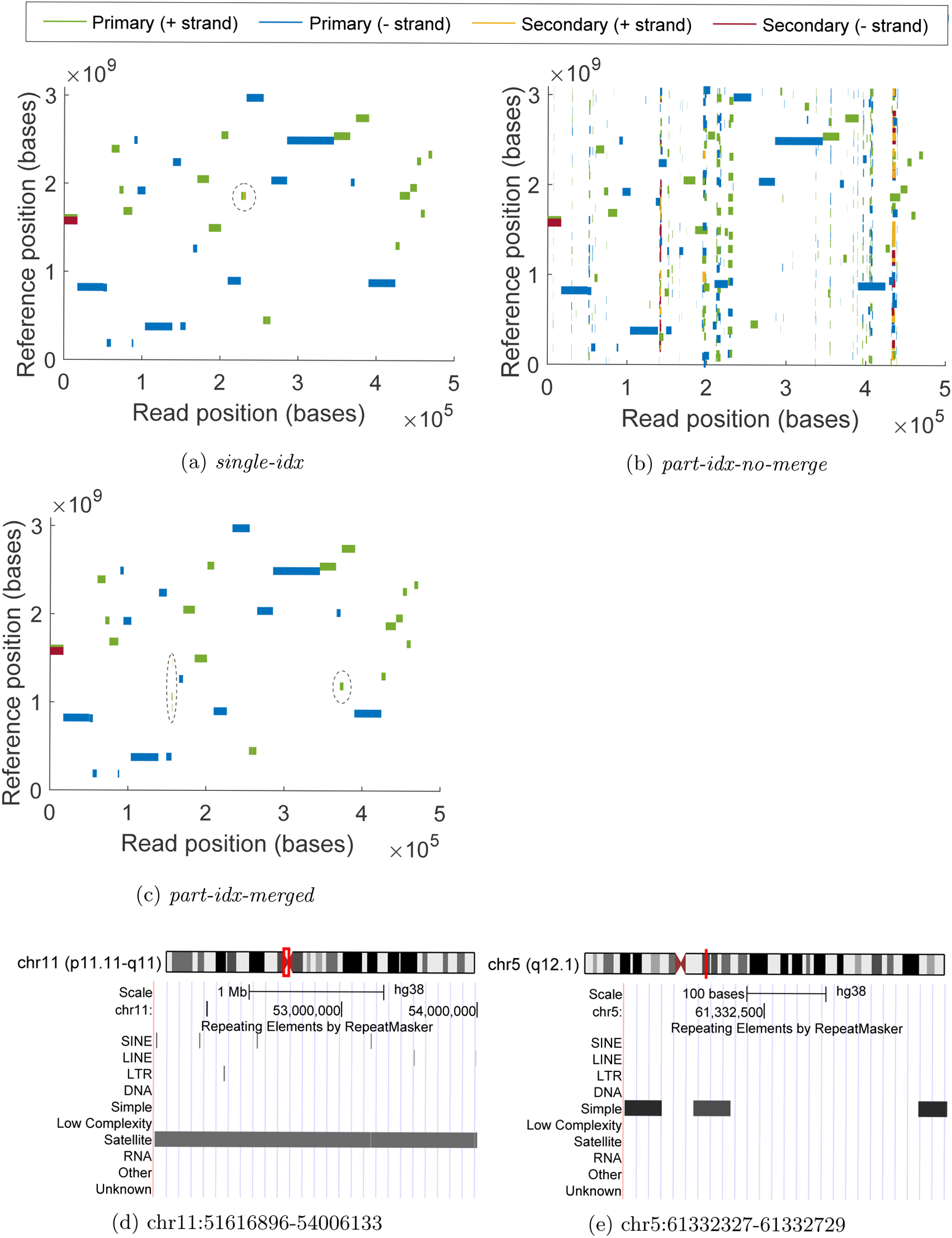
Alignment of an ultra-long nanopore read from a chromothriptic region. Mapping coordinates in the entire human reference genome (y-axis) in function of the position in the read, showing where sub-sequences of the chimeric read map to in the genome for *single-idx* (**a**), *part-idx-no-merge* (**b**), and *part-idx-merged* (**c**). The y-axis begins with chromosome 1 at 0 and ends with chromosome X, Y, and the mitochondria at the top. The length of rectangles along the x-axis are in the correct scale to the length of the read. However, the length along the y-axis are exaggerated to a fixed value so that it is clearly visible. In (**a**) and (**c**), the areas with dotted circles contain the differences between unique mappings for each alignment strategy. Circled regions in (**a**) map to a genomic locus harbouring the satellite repeat displayed in (**d**). Out of the 6 unique mappings in (**c**), the segment with the highest mapping quality (6) maps to the simple repeat containing region displayed in (**e**).

**Figure S1:**
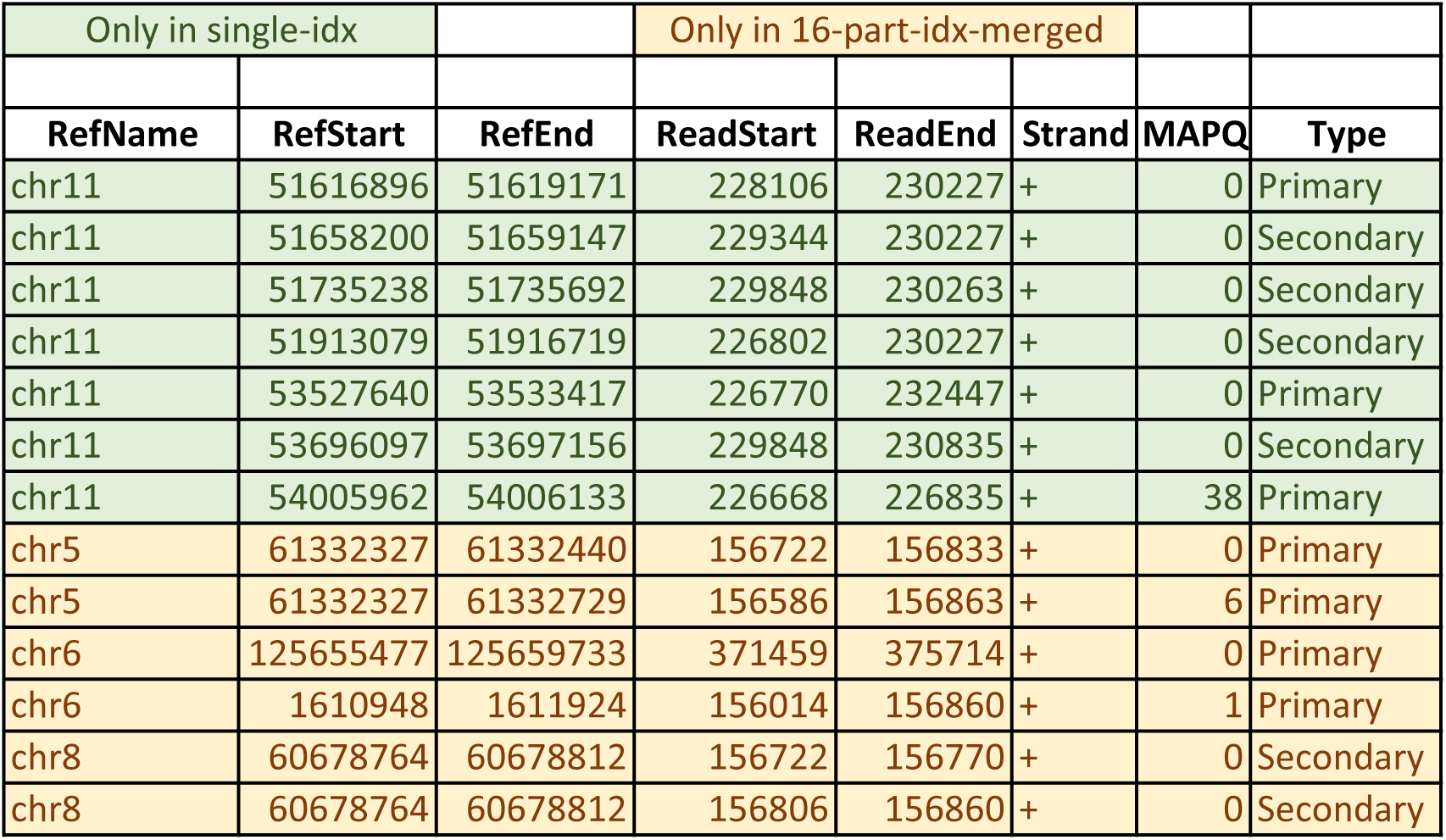
Mappings which are different in single-idx and 16-part-idx-merged. The alignments which were only found in the *single-idx* mapped to locations in the range chr11:51616896-54006133. The ones unique to *16-part-idx-merged* mapped to repetitive regions in chr5, chr6 and chr8. chr5:61332327-61332729, chr6:1610948-1611924 and chr8:60678764-60678812 contained simple repeats. chr6:125,655,477-125,659,733 had simple repeats,SINE repeats and LTR repeats

## Memory usage of partitioned indexes

In addition to the comparable quality of alignments, using a partitioned index yields impressive reductions on peak memory usage (See fig5a). About 7.7GB of memory is required to hold a single reference index, whereas only 1.5GB is needed for a partitioned index with 16 parts. Peak memory usage can be further reduced (to about 1GB for a 16-part index) by balancing chromosomes based on their sizes (see Materials and methods). Hence, using a partitioned index with 16-parts in combination with a batch size between 5-20 Mbases (Minimap2 parameter *K*), aligning long reads to the human genome can be performed on a system with 2GB of RAM without significant loss of accuracy. It is noteworthy to mention that building an index *ab initio* requires slightly more memory than loading a pre-built index (fig5b). This is further remedied by building the index using chromosome balancing, which can be performed *a priori*.

**Figure 5:**
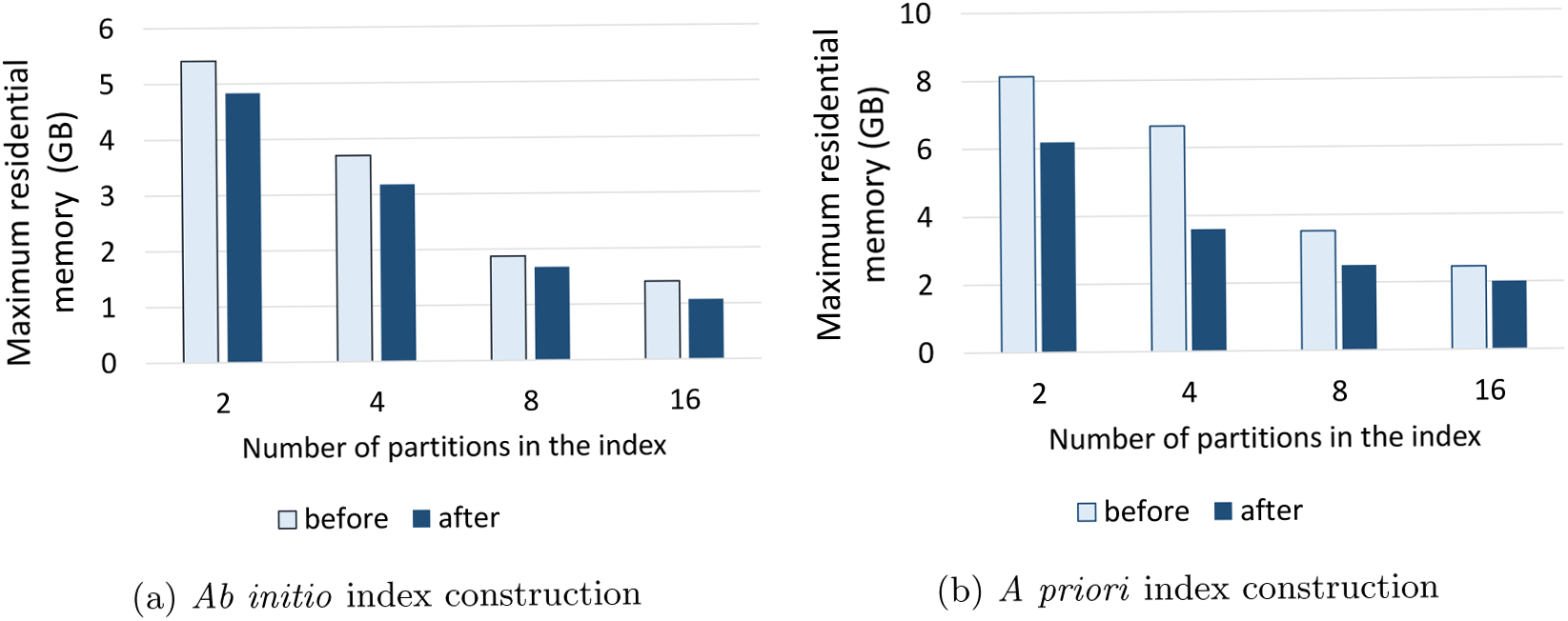
Peak memory usage for a partitioned index. (**a**) Peak memory for holding a pre-built index in function of the amount of partitions. (**b**) Peak memory requirements when loading a pre-built index. Indexing based on the order of chromosomes is referred to as *before* (dark blue bars). Indexing based on the distribution of the sizes of chromosomes is referred to as *after* (light blue bars).

The reduced peak memory usage of a partitioned index comes with an inherent sacrifice in performance in terms of computing speed. We observed that the execution time increased almost linearly with the number of partitions in the index (data not shown), as anticipated.

## Discussion

The partitioned reference approach has been previously used for reducing the memory usage of BWA, a popular short read alignment program [17]. However, the final output is a concatenation of the alignments from all the partitions. We have demonstrated that performing appropriate merging of alignment output is required to eliminate many mapping artefacts, thus improving overall accuracy.

We also showed that *part-idx-merged* can provide a better result than a simple strategy of filtering out results with low mapping quality in *part-idx-no-merge*. This is supported by the results from synthetic reads, where the accuracy of alignments with mapping quality 60 in *part-idx-no-merge* is lower than those from *part-idx-merged*. Furthermore, a simple strategy to remove all short mappings from *part-idx-no-merge* is also less than ideal. In fact, *Paftools* (which was used for evaluating the synthetic read alignments) considers the longest primary mapping when multiple primary mappings exist to assess alignment accuracy.

However, *part-idx-merged* can sometimes generate non-identical alignments to that of *single-idx*. This is like a consequence of slight variations in highly abundant k-mers observed when the index is build. Overall, this affects only a few reads which would nonetheless have low mapping qualities–an issue that has previously been reported by the author and users of Minimap2 (see the public code repository associated with [4]. Further, the reported alignments might differ in long low-complexity regions, as Minimap2 may generate suboptimal alignments in long low-complexity regions (see supplementary materials of [4]).

Although a partitioned index reduces peak memory usage, the runtime is proportionately higher. This is because all the reads should be repeatedly mapped to each partition of the reference. However, this strategy lends itself well to distributed computing, in particular when many smaller, less expensive computing devices are available.

A limitation of this method also lies in the maximal number of partitions an index can be split into, which currently depends on the longest chromosome or contig. We have not yet investigated the impact of splitting chromosomes into fragments, although we anticipate this would not drastically affect results (as exemplified from the chromothriptic read example above). Furthermore, we have not tested the impact of this strategy for RNA sequencing read alignment, which implements different alignment scoring metrics.

In addition to capability of mapping long reads to large genomes on devices with a small memory footprint, our extension to Minimap2 could potentially be useful for the following applications:

- *Mapping to huge reference genome databases*. Meta-genomic databases can be hundreds of gigabytes in size. Hence, holding the index for the whole database would be challenging even for high-specification servers. Especially when multiple species with similar genomes are present, an accurate mapping quality with correct flags, headers, and reduced output file sizes is always appreciated. Alternatively, mapping genome assembly contigs, or a select amount of long reads, to a large public sequence repository (akin to a BLASTN nucleotide databse query) could benefit from our approach. However, the effect of merging output from such large queries has yet to be investigated.
- *Mapping with a lower window size for increased sensitivity*. Minimap2 runs on a default minimiser window size of 10. However, reducing this value improves the mapping sensitivity, but increases the memory consumption. For application where high sensitivity may be preferred, for instance when confronted low coverage sequencing data, our method can be beneficial.

While preparing this manuscript, our method was integrated into the source code of the original Minimap2 software repository. In Minimap2 version 2.12, the option *–split-prefix* can be used to align to a partitioned index. The developer has expanded our implementation to support paired end short reads and multi-threading for the merging process. The original version we implemented for conducting the above experiments is available at https://github.com/hasindu2008/minimap2-arm and can be useful for understanding the underlying algorithm. The partitioned index functionality can be invoked with the option *–multi-prefix*.

## Materials and methods

### Exploration of parameters that affects Memory usage in Minimap2

For measuring peak memory usage and runtime, publicly available NA12878 Nanopore reads [14] were aligned to the human genome reference (GRCh38) with Minimap2 [4]. Peak memory usage and runtime were measured by using the GNU command line *time* utility with the *-v* option. Sensitivity and error rate calculations for different window sizes (Minimap2 parameter *w*) were performed using DNA sequins–chiral, synthetic human genome spike-in standards. The particular Sequins we employed (Deveson et al, under review) were unpublished at the time this manuscript was written, but are conceptually similar to what is reported in [18]. The Sequins were sequenced on ONT MinION sequencers with R9.4.1 flow cells, using LSK108 library preparation kits and the results were base called with ONT’s Albacore sequencing pipeline software version 1.2.6.

Nanopore reads for the sequins were used in this experiment to represent real biological reads. Those nanopore reads were mapped to the reverse human genome, using Minimap2 under the pre-set *map-ont* for different window sizes. However, the exact mapping positions on the reverse human genome for the reads are unknown, given stochastic variations in sequencing (base calling idiosyncrasies, library fragments, etc). Hence, the primary mappings from default window size parameter (*w* = 10) in Minimap2 were used as the truth set. Then, for a particular window size:

- *Mismatching mappings* refer to primary mappings that had different positions to that in the truth set;
- *Missing mappings* refer to primary mappings that were not observed, but were present in the truth set;
- *Extra mappings* refer to primary mappings that were observed in empirical alignments, but were not in the truth set.

The above counts were expressed as a percentage of the total number of reads. The sum of the mismatch and extra mapping percentages were taken as an approximate measure for the error rate. The sensitivity was approximated by subtracting missing mapping percentage from 100.

## Merging of mappings from a partitioned Index

We extended the partitioned index approach of Minimap2 to eliminate alignment artefacts as described below. The index partitioning in Minimap2 is inherited from the first version of Minimap [19]. This feature is for finding long read overlaps to be used with assembly tools such as Miniasm [19]. As overlap computing requires all-vs-all mapping of reads, the index is built for chunks of 4 Gbases (can be overridden with the *-I* argument) at a time, effectively partitioning the alignment index to keep the maximum memory capped at around 27GB. For each part of the index, Minimap2 attempts to map all the reads. The concatenated alignments from all the parts is the final output.

We modified Minimap2 to dump the software’s internal state during the alignment process. The internal state is dumped in binary format to reduce disk usage. The internal state includes: (i) mapped positions, chaining scores and other mapping statistics for each alignment record; (ii) DP score, CIGAR string, and other base-level alignment statistics for each alignment record (when base-level alignment is specified); and (iii) sum of region length of read covered by highly repetitive k-mers for each read (referred to as repeat length). These data form the binary dump files, one for each partition of the index.

When an alignment process has completed, we simultaneously open all the dumped files together with the queried sequence file. For each queried read (or contig), the previously dumped internal states of all alignments for the given read (resulting from all the index partitions) are loaded into memory. If no base-level alignment has been requested, the alignments are sorted based on the chaining score in descending order. Otherwise, the sorting is based on the DP alignment score in descending order. The classification of primary and secondary chains is reiterated as implemented in Minimap2. This corrects the primary and secondary flags in the output. Then, the secondary alignment entries are filtered based on a user requested number of secondary alignments, and the requested minimum primary to secondary score ratio, effectively removing spurious secondary alignments. If a SAM output has been requested, the best primary alignment is retained as the primary alignment and all other primary alignments are classified as supplementary alignments. An unaligned record is printed only if the read is not mapped to any part of the index.

The length of the read covered by repeat regions in the whole genome (repeat length) is required to estimate an ideal mapping quality (MAPQ). We estimate this global repeat length by taking the maximum of the previously dumped repeat lengths (for each partition of the index) for that particular read. The Spearman correlation between this estimated repeat length and the global repeat length is 0.9961. In theory, it would possible to exactly calculate this value by dumping the positions of repeats within the read. However, as the MAPQ is itself an estimation and the accuracy of mappings was adequate in our initial tests, we simply took the maximum. Hence, the computed MAPQ during merging of a partitioned index is not exactly the same as for a single reference index, but very similar overall. This computed MAPQ is more accurate than a MAPQ computed only from the repeat length for a single part of the index.

For space efficiency, dumped data contains numeric identifiers for references (the same used by internal Minimap2 data structures) rather than the reference name. This numeric identifier is determined per index partition and is fixed during the merging using an emulated single reference index. When each index partition is loaded/constructed during mapping, the meta data of reference sequences (sequence name, length, etc) in the index are dumped along with their respective numeric identifiers. At the beginning of merging, these meta-data are reloaded and the numeric identifier is rectified by adding a necessary offset to form an emulated single reference index. This allows generation of a correct SAM header. The numeric identifiers for dumped alignments are also fixed in the same way at the time of loading.

Merging is performed in the order of input read sequences, and mappings for a particular read ID will be adjacent in the output. As the emulated single reference index contains meta-data, and the dumped data are loaded into memory for each read at a time, the memory usage of merging is only a few megabytes. N.B. reads could be batched to improve performance.

## Chromosome balancing

The construction of partitioned indexes in Minimap2 (specified by *-I* option) processes the reference sequences based on their order in the reference. The next part of the index in considered only when the user specified number of bases (4G by default) is exceeded. When building a partitioned index for overlap finding, the parts would be approximately equal in size as the length of the longest read would be in the order of mega bases, at most. However, for reference sequences like the human genome, where the chromosomes are of highly variable lengths, the size of index partitions will be unbalanced. Thus, the largest partition of the index determines the peak memory. We implemented the following strategy to mitigate this effect. First, a command line parameter describing the number of desired partitions is considered. Then, the reference sequences are sorted in descending order based on the sequence length (length without the ambiguous N bases). Next, the sorted list is traversed while putting the current sequence into the bucket with the lowest sum. We output the references sequences belonging to the each bucket in a separate file. Finally we launch the Minimap2 indexer on each file and concatenate the indices.

The extended version of Minimap2 with our modifications can be downloaded from [https://github.com/hasindu2008/minimap2-arm]. Under *misc/idxtools* in this link, we have included the tool that further improves the peak memory for a partitioned index by splitting the reference based on the lengths of the reference sequences, rather than the order of existence in the file.

## Datasets and evaluation methodology

All experiments were performed using the human genome as a reference (GRCh38 with no ALT contigs).

### Synthetic reads

Mapping accuracy was evaluated using synthetic long reads. We generated about 4 million reads pac-Bio reads using PbSim [20] under the Continuous Long Read (long reads with high error rate) mode. Simulated long reads were aligned using Minimap2 with single reference index (*single-idx*), partitioned index without merging (*part-idx-no-merge*) and partitioned index with merging (*part-idx-merged*). Partitioned indices with 2, 4, 8 and 16 parts were tested. For each instance, we evaluated base-level alignments (default SAM output) as well as locus- or block-based alignment (default PAF output with no CIGAR).

To evaluate alignment accuracy, the *Mapeval* utility in *Paftools*–part of the Minimap2 software package–was used with default options. By default, it considers the longest primary alignment for a read. However, *Paftools* assumes that all alignments for a particular read reside contiguously. Hence, for *part-idx-no-merge*, we first sorted the alignments based on the read ID. The output from *Paftools* contains the accumulative mapping error rate and the accumulative number of mapped reads for different mapping quality thresholds [21]. The fraction of mapped reads is taken as a measure of sensitivity.

## Nanopore sequencing data

We could not find a suitable Nanopore simulator. Published Nanopore simulators explored at the time of writing were either dependent on Minimap2 (would cause a bias), did not have models for human genome or unstable. (For instance DeepSimulator [22] and [23] are dependent on Minimap2, SNaReSim [24] code was not available.) Hence we used a a dataset from the publicly available NA12878 sample (rel3-nanopore-wgs-84868110-FAF01132 at [14]). The dataset had 689,781 reads with about 5.5 Gbases. We aligned this dataset to the human genome using a 16-part index with merging (*part-idx-merged*) and without merging (*part-idx-no-merge*) with base-level alignment. Then we compared those outputs by generating some alignment summary metrics with the result from a single reference index (*single-idx*). We initially attempted to perform an extensive comparison using tools such as *CompareSAMs* utility in *Picard* [25] and *qProfiler* utility in *AdamaJava* [26]. They crashed probably because they are designed to be worked with short reads. Hence, we had to limit out test to some simple summary metrics obtained through custom shell scripts.

The ultra-long chromothiptic read was sourced from an unpublished patient-derived dataset generated in house (see [16] for more details on the 778 cancer cell line. The data was generated on a MinION MkI sequencer (MN16218) with MinKNOW version 1.1.17 on a first generation R9 flowcell (MIN105, no spot-on loading, flow cell ID FAD24075) using the SQK-RAD001 library preparation kit from ONT. The raw data for the read was live base called with MinKNOW 1.1.17 and produced an average fastq score of 7.8.

## Conclusion

Aligning long reads generated from third generation high-throughput sequencers to large reference genomes is possible on computers with limited volatile memory. Parameter optimisation alone cannot substantially reduce memory usage without considerably sacrificing alignment quality. Partitioning an alignment index, saving the internal state, and merging the output *a posteriori* substantially reduces memory usage. This strategy reduces the memory requirements for aligning nanopore reads to the human reference genome from 11GB to less than 2GB, with minimal impact on accuracy.

## Acknowledgements

We thank Heng Li, the author of Minimap2 for assisting us in understanding the code, providing us with valuable insights and suggestions through Github issues, and integrating our method into his original software. We also thank Prof David Thomas, Prof Tony Papenfuss, Prof Vanessa Hayes and, in particular, Ruth Lyons for their assistance with generating sequencing data from which the (once record-holding) ultra-long chromothriptic nanopore read was sourced.

---

**Table S1:** Memory usage of Minimap2 for default parameters Minimap2 was run with default parameters. Pre-set profiles *map-pb* and *map-ont* were used for PacBio and Oxford Nanopore respectively. The peak memory usage for each event has been reported. Index construction refers to the building of the index and then dumping to a file. Index residence is the memory required only for the index to reside in memory.

|  | PacBio | Oxford nanopore |
| --- | --- | --- |
| Index construction | 8.67GB | 11.32GB |
| Index residence | 6.33GB | 7.71GB |
| Mapping with base-level alignment (SAM output) | 8.56GB | 11.30GB |
| Mapping without base-level alignment (PAF output) | 7.14GB | 8.80GB |

